# The validity and consistency of continuous joystick response in perceptual decision-making

**DOI:** 10.1101/501536

**Authors:** Maciej J. Szul, Aline Bompas, Petroc Sumner, Jiaxiang Zhang

## Abstract

A computer joystick is an efficient and cost-effective response device for recording continuous movements in psychological experiments. Movement trajectories and other measures from continuous responses have expanded the insights gained from discrete responses (e.g. button presses) by providing unique information on how cognitive processes unfold over time. However, few studies have evaluated the validity of joystick responses with reference to conventional key presses, and response modality can affect cognitive processes. Here, we systematically compared human participants’ behavioural performance of perceptual decision-making when they responded with either joystick movements or key presses in a four-alternative motion discrimination task. We found evidence that the response modality did not affect raw behavioural measures including decision accuracy and mean reaction time (RT) at the group level. Furthermore, to compare the underlying decision processes between the two response modalities, we fitted a drift-diffusion model of decision-making to individual participant’s behavioural data. Bayesian analyses of the model parameters showed no evidence that switching from key presses to continuous joystick movements modulated the decision-making process. These results supported continuous joystick actions as a valid apparatus for continuous movements, although we highlighted the need for caution when conducting experiments with continuous movement responses.

## Introduction

Discrete key presses on a keyboard or button box have been the long-standing response modality in computer-based experiments in psychology, from which on/off responses and response time (RT) are commonly measured. Developments in computers and electronics technology have improved the accessibility of other devices that are capable of recording continuous responses, e.g., joystick, computer mouse, motion sensor and robotic arm (Koop & Johnson, 2011; O’Hora, Dale, Piiroinen, & Connolly, 2013). In addition to the standard behavioural measures available from key presses, continuous responses enable further inferences from movement trajectories. However, to utilize the full capacity of continuous response recording, we need to ensure that experimental results from these devices are consistent with, or generalizable to, the findings from conventional response modalities such as key presses. The current study addressed this issue by comparing the behavioural performance between joystick movements and key presses in a perceptual decision-making task. Using computational modelling of behavioural data, we further compared the decision-making processes from the two response modalities.

### Continuous and discrete responses in experimental psychology

Continuous responses can offer theoretical and practical advantages in experiments. First, although a discrete response is consistent with the assumption of sequential stages of cognition and motor outputs, a growing number of studies suggested a continuous and parallel flow of information between brain systems involved in sensory, cognitive and motor processes (Cisek & Kalaska, 2005; Spivey, Grosjean, & Knoblich, 2005). Continuous responses can capture the dynamics of these multiple mental processes, as well as the transitions between them (Resulaj, Kiani, Wolpert, & Shadlen, 2009). Second, in experiments involving clinical populations, it can be difficult for patients to make discrete responses accurately on a keyboard, especially in patients with dementia or parkinsonism. Patients with motor function impairments (e.g., tremor, apraxia or loss of dexterity) often omit button presses, press the button too early or too late, press wrong buttons accidentally or are confused with response-button mapping. This limitation may result in a significant amount of experiment data being rejected in some studies (Wessel, Verleger, Nazarenus, Vieregge, & Kömpf, 1994), while continuousresponses with natural movements can be well tolerated in patients (Limousin et al., 1997; Strafella, Dagher, & Sadikot, 2003)

The trajectories of continuous movements contain rich spatiotemporal information of the action, and provide unique insights into how cognitive processes unfold in time (Freeman, Dale, & Farmer, 2011; Song & Nakayama, 2009). In continuous reaching, movement trajectories showed that human participants can initiate a reaching action prior to when the target becomes fully available, and select from competing action plans at a later stage (e.g. Chapman et al., 2010; Gallivan & Chapman, 2014). In perceptual decision-making, movement trajectories from joysticks and other similar devices have been successfully used to investigate the cognitive processes underlying changes of mind (Resulaj et al., 2009), error correction (Acerbi, Vijayakumar, & Wolpert, 2017) and subjective confidence (Berg et al., 2016) that are otherwise difficult to study with key presses.

### A comparison between response modalities

To extend currently available experimental findings to otherdevices, it is necessary to assess the consistency of performance between response modalities. More importantly, characterising the consistency between response modalities may help us understand the interdependence of cognitive processes and motor systems. For example, in decision-making tasks, comparisons between saccadic eye movements and manual responses has suggested a domain general decision mechanism regardless of response modality (Gomez, Ratcliff, & Childers, 2015; Ho, Brown, & Serences, 2009), and the apparent difference in response speed is accounted for by the neuroanatomical distinctions in saccadic and manual networks (Bompas, Hedge, & Sumner, 2017).

The current study aimed to examine the validity and consistency of continuous joystick responses versus discrete button presses in perceptual decision-making. Participants performed a four-alternative motion discrimination task (Churchland, Kiani, & Shadlen, 2008) with two levels of perceptual difficulty. The task was to indicate the coherent motion direction from random dot kinematogram, a standard psychophysical stimulus for visual perceptual decision (Fredericksen, Verstraten, & Van De Grind, 1994; Lappin & Bell, 1976; Pilly & Seitz, 2009; Ramachandran & Anstis, 1983; Watamaniuk, Sekuler, & Williams, 1989). In two counterbalanced sessions, the participants indicated their decisions with either joystick movements or key presses. The joystick response was to move the lever from its neutral position towards one of the four cardinal directions, aligned to the coherent motion direction, and the corresponding key press was one of the four arrow keys on the keyboard. We compared raw behavioural performance (decision accuracy and mean RT) between the two response modalities and between the two levels of task difficulty. From continuous movement trajectories, we also examined whether joystick-specific measures were consistent between movement directions (i.e., trajectory length, peak velocity and acceleration time).

To assess whether the response modality affected the decision-making process, we fitted a drift-diffusion model (DDM) (Gold & Shadlen, 2007; Ratcliff, Smith, Brown, & McKoon, 2016) to individual participant’s behavioural data and compared model parameters derived from the joystick and keyboard sessions. The DDM belongs to a family of sequential sampling models of reaction time. These models assume that the decision process is governed by the accumulation of noisy sensory evidence over time until a threshold is reached (Bogacz, Brown, Moehlis, Holmes, & Cohen, 2006; Ratcliff & Smith, 2004), consistent with the electrophysiological (Britten, Shadlen, Newsome, & Movshon, 1992; Churchland et al., 2008; Hanks, Kiani, & Shadlen, 2014; Huk & Shadlen, 2005; Shadlen & Newsome, 2001) and neuroimaging (Heekeren, Marrett, & Ungerleider, 2008; Ho, Brown, & Serences, 2009; Zhang, Hughes, & Rowe, 2012) evidence on the identification of neural accumulators in the frontoparietal cortex. The current study used the DDM to decompose the observed RT distributions and accuracy into three main model components: decision threshold for the amount of evidence needed prior to a decision, drift rate for the speed of evidence accumulation, and non-decision time to account for the latencies of stimulus encoding and action initiation (Karahan, Costigan, Graham, Lawrence, & Zhang, 2019; Ratcliff & McKoon, 2008; Wagenmakers, 2009; Zhang, 2012). The latter parameter is of interest, because one may expect a difference in the latency distribution of action initiation between joystick movements and key presses.

Our findings demonstrated that when human participants used ballistic movements to respond with a joystick, their behavioural performance was modulated by task difficulty and similar to that from key presses during the same perceptual task. Further computational modelling analysis showed no evidence of a change in any model parameter when switching between response modalities. As such, we concluded that joystick movement is a valid response modality for extending discrete actions to continuous behaviour in psychological experiments, although participants may exhibit differences in movement trajectory measures towards different directions.

## Method

### Participants

Twenty-one participants (7 males, aged range 18-24 years; M = 20.43 years, SD = 2.91 years) took part in the study following written informed consent. All but three were right-handed. All the participants had normal or corrected-to-normal vision, and none reported a history of motor impairments or neurological disorders. The study was approved by the Cardiff University School of Psychology Ethics Committee.

### Apparatus

The experiment was conducted in a behavioural testing room with dimmed light. Stimuli were displayed on a 22-inch CRT monitor with 1600×1200 pixels resolution and 85 Hz refresh rate. A chin rest was used to maintain the viewing distance and position. A joystick (Extreme 3D Pro Precision, Logitech International S.A., Switzerland) was used to record movement trajectories at 85 Hz in the joystick session. The experimental setup for joystick and keyboard sessions was illustrated in Supplementary Figure 1. The joystick handle could move nearly freely, with little resistance from its neutral position within the 20% movement radius. Beyond the 20% radius, the resistance during joystick movement was approximately constant. A standard PC keyboard was used to record key presses. The experiment was written using PsychoPy 1.85.4 library (Peirce, 2008).

### Stimuli

In both the joystick and keyboard sessions, a random-dot kinematogram was displayed within a central invisible circular aperture of 14.22° diameter (visual angle). White dots were presented on a black background (100% contrast) with a dot density of 27.77 dots per deg^2^ per second and a dot size of 0.14°. Similar to previous studies (Britten et al., 1992; Pilly & Seitz, 2009; Roitman & Shadlen, 2002; Shadlen & Newsome, 2001; Zhang & Rowe, 2014), we introduced coherent motion information by interleaving three uncorrelated sequences of dot positions across frames at 85 Hz. In each frame, a fixed proportion (i.e., the motion coherence) of dots was replotted at an appropriate spatial displacement in the direction of the coherent motion (51.195°/s velocity), relative to their positions three frames earlier, and the rest of the dots were presented at random locations within the aperture. Signal dots had a maximum lifetime of three frames, after which they were reassigned to random positions. The coherent motion direction in each trial was set in one of the four cardinal directions (0°, 90°, 180° or 270°).

### Task and procedure

Each participant took part in two experimental sessions using keyboard or joystick as a response modality. The order of response modality was counterbalanced across participants. In both sessions, participants performed a four-alternative motion discrimination task, indicating the coherent motion direction from four possible choices (0°, 90°, 180° or 270°). Each session comprised 960 trials, which were divided into 8 blocks of 120 trials. Each block had 15 repetitions of each of the four motion directions and two difficulty conditions. The motion coherence was set to 10% in the “Difficult” condition and 20% in the “Easy” condition. Feedback on the mean decision accuracy was provided after each block. The order of the conditions was pseudo-randomized across sessions and participants, ensuring that the same direction and difficulty condition did not occur in four consecutive trials. In the keyboard session, the participants responded with four arrow keys corresponding to the coherent motion direction (right - 0°, up - 90°, left - 180° and down - 270°). In the joystick session, the participants were instructed to indicate the motion direction with an appropriate joystick movement from the joystick’s central position towards one of the four edges (right - 0°, up - 90°, left - 180° and down - 270°).

Every trial started with a 400 ms fixation period (Figure 1a). The random dot kinematogram appeared after the fixation period for a maximum of 3000 ms or until response. In the keyboard session, stimuli disappeared after a button press. In joystick condition, stimuli disappeared when the participants stopped joystick movement. The chosen stopping rule was when the joystick position did not change in the last four sampling points, and its position was outside of the 20% motion radius. After response, a blank screen was presented as the intertrial interval, with a duration uniformly randomized between 1000 and 1400 ms.

**Figure 1.**
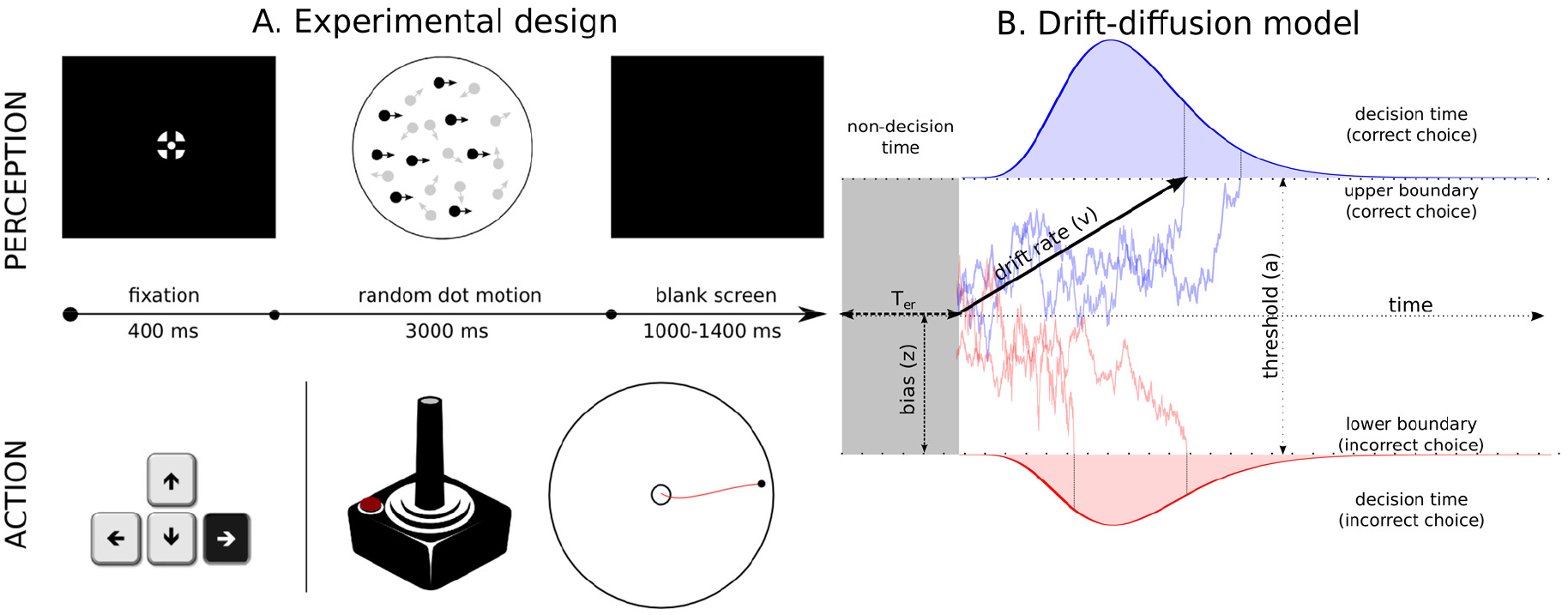
Behavioural paradigm and the drift-diffusion model (DDM). (**A**) The structure of a single trial of the experiment. A fixation screen was presented for 400 ms, after which the random-dot kinematogram was presented for a maximum of 3000 ms or until response. The inter-trial interval was randomised between 1000 and 1400 ms. Participants were instructed to indicate the direction of the coherent motion direction (0°, 90°, 180° or 270°) using joystick or keyboard in two counterbalanced sessions. (**B**) The drift-diffusion model and examples of evidence accumulation trajectories. The parameter (*a*) indicates the distance between the correct and incorrect decision thresholds. The drift rate (*v*) represents the speed of evidence accumulation and its magnitude is determined by the quality of the evidence. A positive *v* indicates that, on average, the accumulation of sensory evidence is towards the correct decision threshold. The starting point (*z*) represents the response bias towards one of the two thresholds. The non-decision time (*T*_*er*_) represents the latencies of non-decision processes, which is illustrated besides the decision time distribution in the figure. The diffusion process starts at the starting point (*z*) until the accumulated evidence reaches one of the two thresholds. If the accumulated evidence reaches the correct (upper) threshold (blue trajectories), the model predicts a correct response. Because of noise, the accumulated evidence may reach the incorrect (lower) threshold (red trajectories) and the model predicts an incorrect response. The predicted single-trial RT is the sum of the duration of the evidence accumulation (decision time) and the non-decision time *T*_*er*_.

The response time (RT) in the keyboard session was defined as the latency between the onset of random-dot kinematogram and the time of key press. In the joystick session, the RT was defined as the duration between the onset of the random-dot kinematogram and the first time when the joystick’s position left the 20% movement radius from its neutral position. It coincided with the first noticeable increase in the velocity of the movement from the stimulus onset. Participants’ choice in the joystick session was one of the four cardinal directions (i.e., 0°, 90°, 180° and 270°) closest to the last position of the joystick.

### Drift-diffusion model (DDM) analysis

We fitted the DDM to each participant’s response time distributions and accuracy. The DDM decomposes the behavioural data into four key model parameters (Ratcliff & McKoon, 2008). (1) The decision threshold (*a*) denotes the distance between the two decision boundaries. (2) The mean drift rate (*v*) denotes the strength of sensory information. (3) The starting point (*z*) denotes the response bias towards one of the two alternatives. (4) The non-decision time (*T*_*er*_) denotes the latencies of stimulus encoding and response initiation. In addition, the DDM can be extended to include trial-by-trial variability in drift rate *s*_*v*_ and non-decision time *s*_*t*_, which improves model fit to the data (Ratcliff & McKoon, 2008). The DDM predicts the decision time as the duration of the accumulation process and the observed RT as the sum of the decision time and *T*_*er*_ (Figure 1B).

Similar to previous studies (Churchland et al., 2008), we simplified the four-alternative forced choice task in the current study to a binary decision problem for model fitting. This was achieved by separately grouping trials with correct responses and trials with incorrect responses. The behavioural task was then reduced to a binary choice between a correct and an incorrect alternative. We used the hierarchical drift-diffusion model (HDDM) toolbox to fit the behavioural data (Wiecki, Sofer, & Frank, 2013). The HDDM implemented a hierarchical Bayesian model (Vandekerckhove, Tuerlinckx, & Lee, 2011) for estimating the DDM parameters, which assumes that the model parameters for individual participants are sampled from group-level distributions at a higher hierarchy. Given the observed experimental data, the HDDM used Markov chain Monte Carlo (MCMC) approaches to estimate the joint posterior distribution of all individual- and group-level parameters. The posterior parameter distributions can be used directly for Bayesian inference (Gelman et al., 2014), and this Bayesian approach has been shown to be robust in recovering model parameters when limited data are available (Ratcliff & Childers, 2015; Wiecki et al., 2013; Zhang et al., 2016).

We applied a few constraints to the model parameters based on our task design. First, we allowed all the model parameters (*a, v, Ter, s*_*v*_, and *s*_*t*_) to vary between the two response modalities. Second, the mean drift rate *v* was further allowed to vary between task difficulties (easy, difficult) and correct directions (up, down, left and right). Third, the starting point *z* was fixed at 0.5, suggesting that there was no bias towards the two decision boundaries and the equal amount of evidence was required for a correct and incorrect decision. This was because the participants did not have *a priori* knowledge about the correct alternative at the beginning of each trial.

We generated 15,000 samples from the joint posterior distribution of all model parameters by using MCMC sampling (Gamerman & Lopes, 2006). The initial 7,000 samples were discarded as burn-in for stable posterior estimates. Geweke diagnostic (Cowles & Carlin, 1996) and autocorrelation were used to assess the convergence of the Markov chains in the last 8,000 samples. All parameter estimates were converged after 15,000 samples.

### Data analysis

First, we used both Bayesian and frequentist repeated-measures ANOVA to make inferences on behavioural measures (JASP Team, 2018). For frequentist ANOVAs, Greenhouse-Geisser correction was applied when the assumption of sphericity was violated. For Bayesian ANOVAs, we followed the standard heuristic to characterize the strength of evidence based on the Bayes factor (BF_10_) (Wagenmakers, Lee, Lodewyckx, & Iverson, 2008), which can provide evidence supporting either null (BF_10_<1) or alternative (BF_10_>1) hypotheses. A BF_10_ between [1, 3] (or [0, 1/3]) suggests weak evidence for the alternative (or null) hypothesis. A BF_10_ between [3, 10] (or [1/10, 1/3]) suggests moderate or compelling evidence for the alternative (or null) hypothesis. A BF_10_ larger than 10 (or smaller than 1/10), suggests strong evidence for the alternative (or null) hypothesis.

Second, to quantify the difference of RT distributions between response modalities, we used the Kolmogorov-Smirnov test (Pratt & Gibbons, 1981), a non-parametric statistical measure of difference between two one-dimensional empirical distributions.

Third, to compare a fitted DDM parameter between two conditions (e.g., between response modalities or between task difficulties), we used Bayesian hypothesis testing (Bayarri & Berger, 2004; Gelman et al., 2014; Kruschke, 2015; Lindley, 1965) to make inferences from the posterior parameter distributions, under the null hypothesis that the parameter value is equal between the two conditions.

More specifically, we first calculated the distribution of the parameter difference from the two MCMC chains of the two conditions, and we obtained the 95% highest density interval (HDI) of that difference distribution between the two conditions. We then set a region of practical equivalence (ROPE) around the null value (i.e., 0 for the null hypothesis), which encloses the values of the posterior difference that are deemed to be negligible from the null value 0 (Kruschke, 2013). In each Bayesian inference, the ROPE was set empirically from the two MCMC chains of the two conditions under comparison. For each of the two conditions, we calculated the 95% HDI of the difference distribution between odd and even samples from that condition’s MCMC chain. This 95% HDI from a single MCMC chain can be considered as negligible values around the null, because posterior samples from different portions of the same chain are representative values of the same parameter. That is, we accepted that the null hypothesis is true when comparing the difference between odd and even samples from the same MCMC chain. The ROPE was then set to the widest boundaries of the two 95% HDIs of the two conditions.

From the 95% HDI of the difference distribution and the ROPE, a Bayesian *P*-value was calculated. To avoid confusion, we used *p* to refer to classical frequentist *p*-values, and *P*_*p*_|D to refer to Bayesian *P*-values based on posterior parameter distributions. If ROPE is completely contained within 95% HDI, *P*_*p*_|D = 1 and we accept the null hypothesis (i.e., the parameter values are equal between the two conditions). If ROPE is completely outside 95% HDI, *P*_*p*_|D = 0 and we reject the null hypothesis (i.e., the parameter values differ between the two conditions). If ROPE and 95% HDI partially overlap, *P*_*p*_|D equals to the proportion of the 95% HDI that falls within the ROPE, which indicates the probability that the parameter value is *practically* equivalent between the two conditions (Kruschke & Liddell, 2018).

## Results

### Behavioural results

The behavioural performance of the four-alternative motion discrimination task was quantified by accuracy (proportion of correct responses, Figure 2A) and mean reaction time (RT, Figure 2B). We compared the behavioural performance between response modalities (joystick or keyboard), task difficulties (easy or difficult) and motion directions (up, down, left or right) using three-way Bayesian and frequentist repeated-measure ANOVAs. Across the two response modalities, participants showed decreased accuracy (BF_10_ = 5.112 × 10^30^; *F*(1,20) = 292.709, *p* < 0.001) and increased mean RT (BF_10_ = 1.458 × 10^18^; *F*(1,20) = 63.163, *p* < 0.001) in the more difficult condition. There was compelling evidence against the main effect of response modality on accuracy (BF_10_ = 0.124; *F*(1,20) = 0.083, *p* = 0.776) and weak evidence against the main effect of response modality on mean RT (BF_10_ = 0.560; *F*(1,20) = 0.495, *p* = 0.490). These results indicated similar behavioural performance between joystick and keyboard responses.

**Figure 2.**
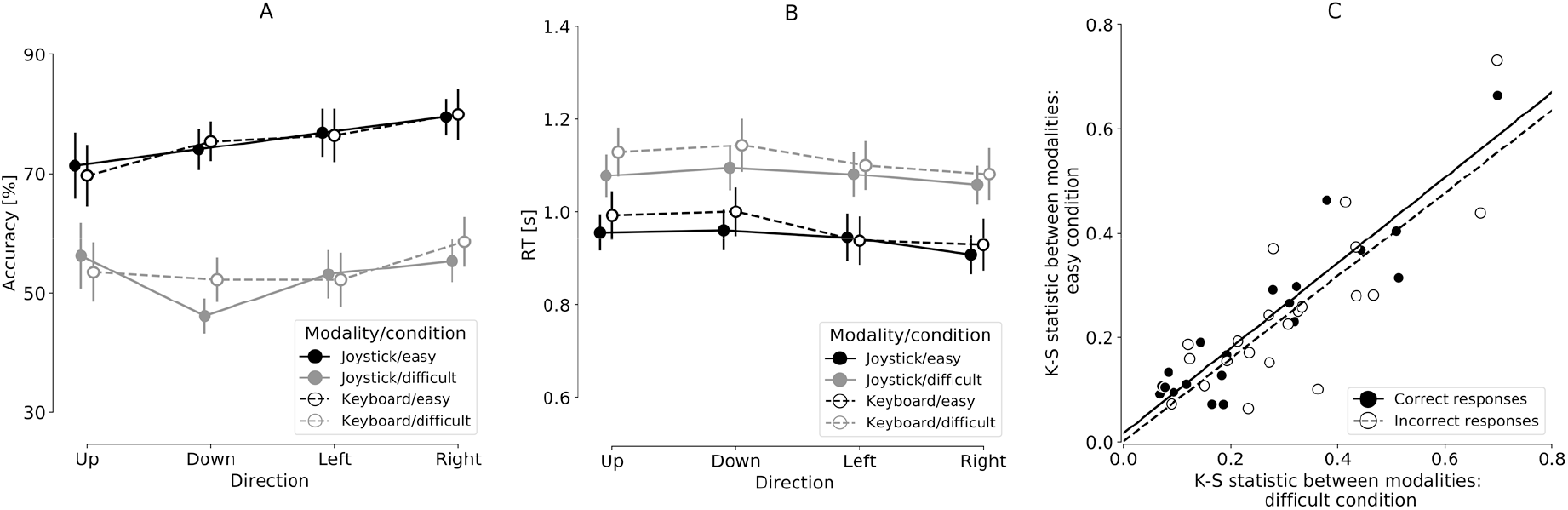
Behavioural results in joystick and keyboard sessions. (**A**) Average decision accuracy (proportion of correct) across participants. Error bars denote standard errors of the means. (**B**) Average mean RT across participants. Error bars denote standard errors of the means. (**C**) The Kolmogorov-Smirnov (K-S) statistics when comparing the RT distributions between response modalities. The scatter plot showed the K-S statistics in the difficult condition as a function of the that in the easy condition. Each data point represents the correct (filled data point) or incorrect (open data point) trials of one participant. Linear regression lines were illustrated for correct (solid line) and incorrect (dashed line) trials.

When comparing the behavioural performance between motion directions, there was compelling evidence against the main effect on accuracy (BF_10_ = 0.185; *F*(2.248, 44.961) = 0.107, *p* = 0.357). On mean RT, the frequentist ANOVA suggested a significant main effect of motion direction (*F*(2.853, 57.052) = 3.021, *p* = 0.039), but this results was supported by neither post-hoc tests (*p*>0.139 in all post-hoc comparisons, Bonferroni corrected) or Bayesian ANOVA (BF_10_ = 0.305). Furthermore, there was a significant interaction on accuracy between task difficulty and motion direction (*F*(2.586, 51.718) = 6.317, *p* = 0.002), although this was again not supported by Bayesian analysis (BF_10_ = 0.299). There was evidence against all the other interactions on accuracy (BF_10_ < 0.179; *p* > 0.228) and mean RT (BF_10_ < 0.199; *p* > 0.083).

The results above suggested no systematic bias at the group level when comparing responses from a joystick and a keyboard. However, the consistency of behavioural performance between response modalities could vary between participants. For experiments with multiple response modalities, the researcher may want to confirm whether the consistency between response modalities is maintained across experimental conditions. This would allow, for example, a pre-screening procedure to identify participants with high response consistency to be recruited for further experiments. Here, we used Kolmogorov-Smirnov (K-S) statistics to quantify the difference of individual participant’s RT distributions between the joystick and keyboard sessions in each difficulty condition, separately for correct and incorrect trials. There was strong evidence of a positive correlation between the K-S statistics of the easy and difficult conditions (correct trials: BF_10_ = 3.647 × 10^6^, *R*= 0.92, *p* < 0.001; incorrect trials: BF_10_ = 4526.00, *R* = 0.82, *p* < 0.001) (Figure 2C). Therefore, the difference in behavioural performance between response modalities was consistent within participants across difficulty levels.

### Hierarchical drift-diffusion model analyses

To compare the underlying decision-making process between joystick and keyboard responses, we simplified the four-alternative motion discrimination task to a binary decision task (Churchland et al., 2008; see also “Drift-diffusion model” section) and fitted the drift-diffusion model (DDM) to the behavioural data using the hierarchical DDM (HDDM) toolbox (Wiecki et al., 2013). The DDM decomposed individual participant’s behavioural data into model parameters of latent psychological processes, and the HDDM toolbox allowed to estimate the joint posterior estimates of model parameters using hierarchical Bayesian approaches. To evaluate the model fit, we generated model predictions by simulations with the posterior estimates of the model parameters. There was a good agreement between the observed data and the model simulations across response modalities, task difficulties and motion directions (Figure 3).

**Figure 3.**
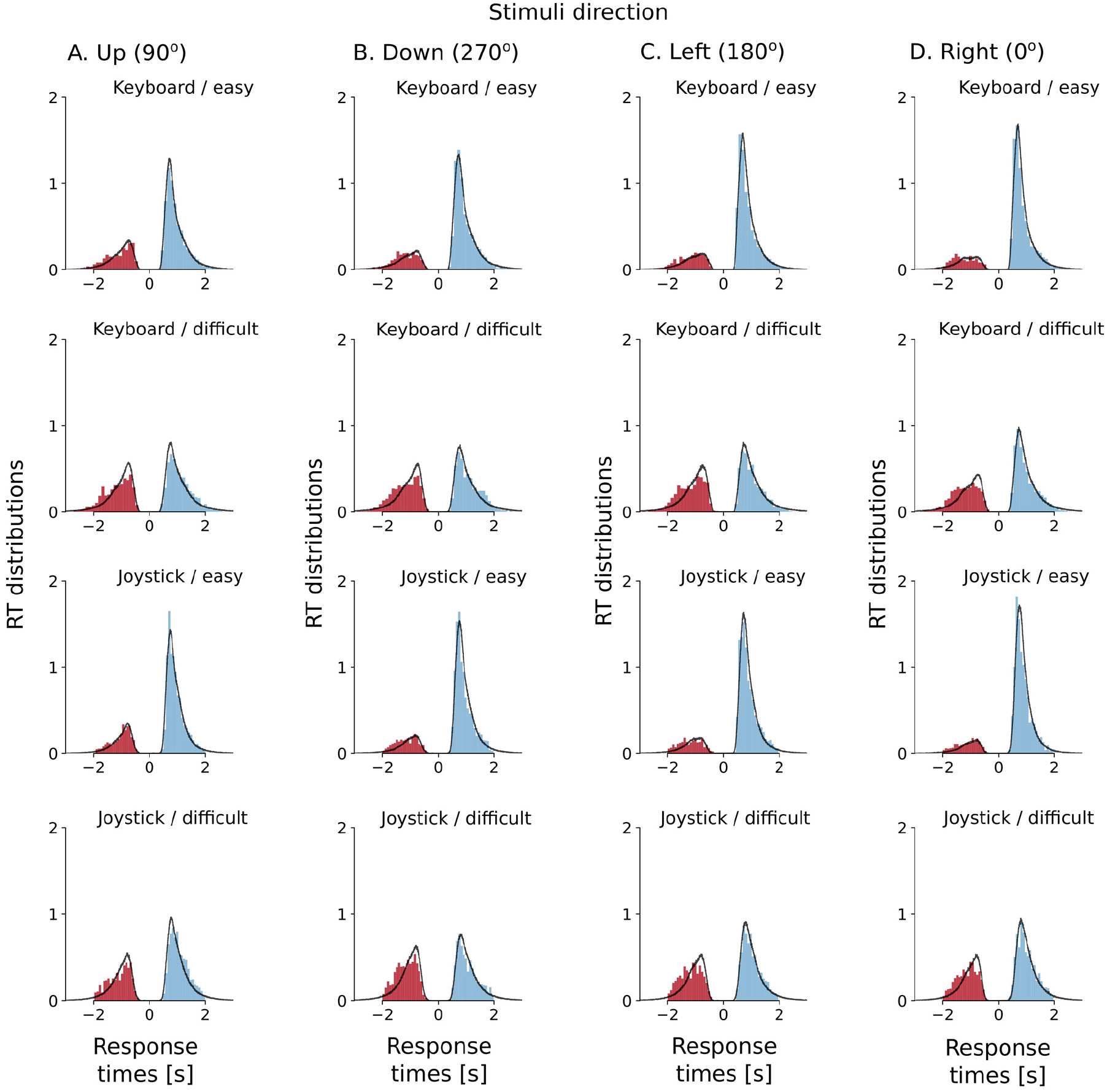
Posterior predictive RT distributions from the fitted DDM. Each panel shows the normalized histograms of the observed data (blue bars – correct responses, red bars – incorrect responses) and the model prediction (black lines) across participants. The RT distribution along the positive x-axis is from correct responses, and the areas under the curve on the positive x-axis corresponds to the observed and predicted accuracy. The RT distribution along the negative x-axis is from error responses, and the areas under the curve on the negative x-axis corresponds to the observed and predicted error. The posterior predictions of the model were generated by averaging 500 simulations of the same amount of model predicted data as observed in the experiment using posterior parameter estimates.

With no *a priori* knowledge on the effect of response modality on the decision-making process, we allowed all model parameters to vary between joystick and keyboard responses: the boundary separation *a*, the mean drift rate *v*, the mean non-decision time *T*_*er*_, the trial-by-trial variability of drift rate *s*_*v*_, and the trial-by-trial variability of non-decision time *s*_*t*_ (Table 1). The mean drift rate was further allowed to vary between task difficulties and motion directions. We performed Bayesian hypothesis testing on the posterior parameter estimates between response modalities (Bayarri & Berger, 2004; Gelman et al., 2014; Kruschke, 2015; Lindley, 1965). This analysis yielded 95% HDI of the parameter difference between the joystick and keyboard sessions, as well as Bayesian *P*-values *P*_*P*|*D*_ (see “Data analysis” section for details).

**Table 1.**
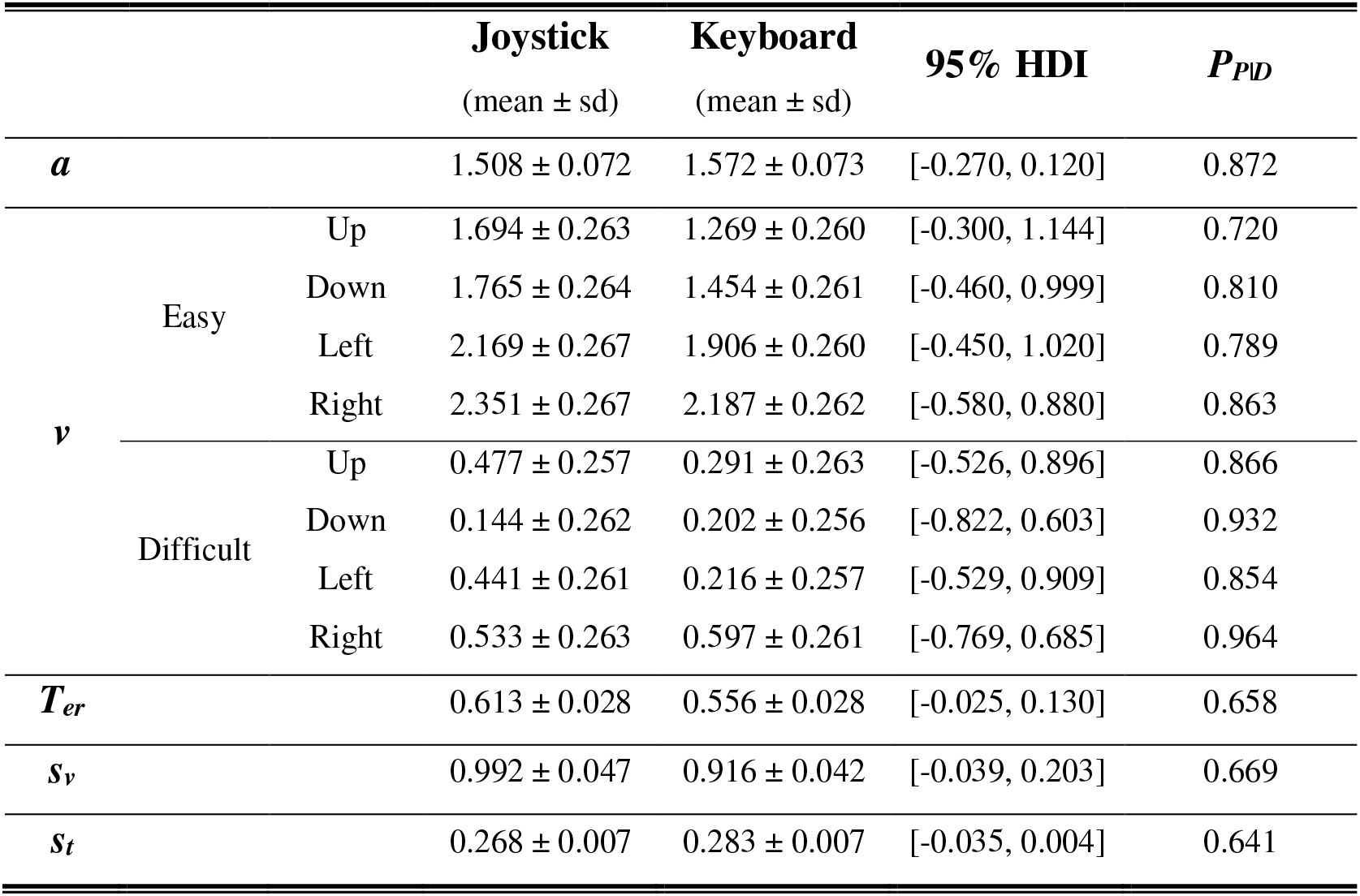
Posterior estimates of the hierarchical drift-diffusion model parameters (decision threshold *a*, mean drift rate *v*, non-decision time *T*_*er*_, trial-by-trial drift rate variability *s*_*v*_, trial-by trial non-decision time variability *s*_*t*_). The first two data columns showed the posterior means and standard deviations of the parameters in the joystick and keyboard sessions. 95% HDI denoted the 95% highest density intervals for the parameter difference between the joystick and keyboard sessions. *P*_*P*|*D*_ denoted the Bayesian *P*-value for the parameter difference being equal between response modalities.

For all the model parameters, we could not reject the null hypothesis that the posterior parameter estimates are practically equal between the joystick and keyboard sessions. The *P*_*P*|*D*_, which quantifies the probability that the model parameter is practically equal between the two conditions, ranged from 0.641 to 0.964 (Table 1). Therefore, there was no evidence to support that switching from keyboard to joystick altered the decision-making process. Next, because the mean drift rate is often assumed to increase with decreased task difficulty (Ratcliff & McKoon, 2008), we compared the drift rate averaged from the joystick and keyboard sessions between easy and difficult conditions. As expected, the drift rate was larger in the easy compared with the difficult condition in all motion directions (up: 95% HDI = [0.589, 1.613], *P*_*P*|*D*_=0; down: 95% HDI = [0.930, 1.958], *P*_*P*|*D*_=0; left: 95% HDI = [1.204, 2.227], *P*_*P*|*D*_=0; right: 95% HDI = [1.185, 2.214], *P*_*P*|*D*_=0).

### Additional measures from joystick trajectories

In the joystick session, the participants’ movement trajectories were close to the four cardinal directions (Figure 4A). Continuous movements with the joystick enabled to acquire additional single trial behavioural measures beyond that possible from simple key presses. We examined three such measures: peak velocity (Figure 4B), acceleration time (Figure 4C) and trajectory length (Figure 4D). These additional joystick measures were subsequent to accuracy and RT. In the current study, we did not expect them to have critical influence on the two primary behavioural measures. Hence our analyses were focused on the effects of movement direction and task difficulty on the trajectory measures. However, we acknowledged that, in experiments with more complex movement trajectories, decisions may be more directly coupled to continuous motor responses (Song & Nakayama, 2009).

**Figure 4.**
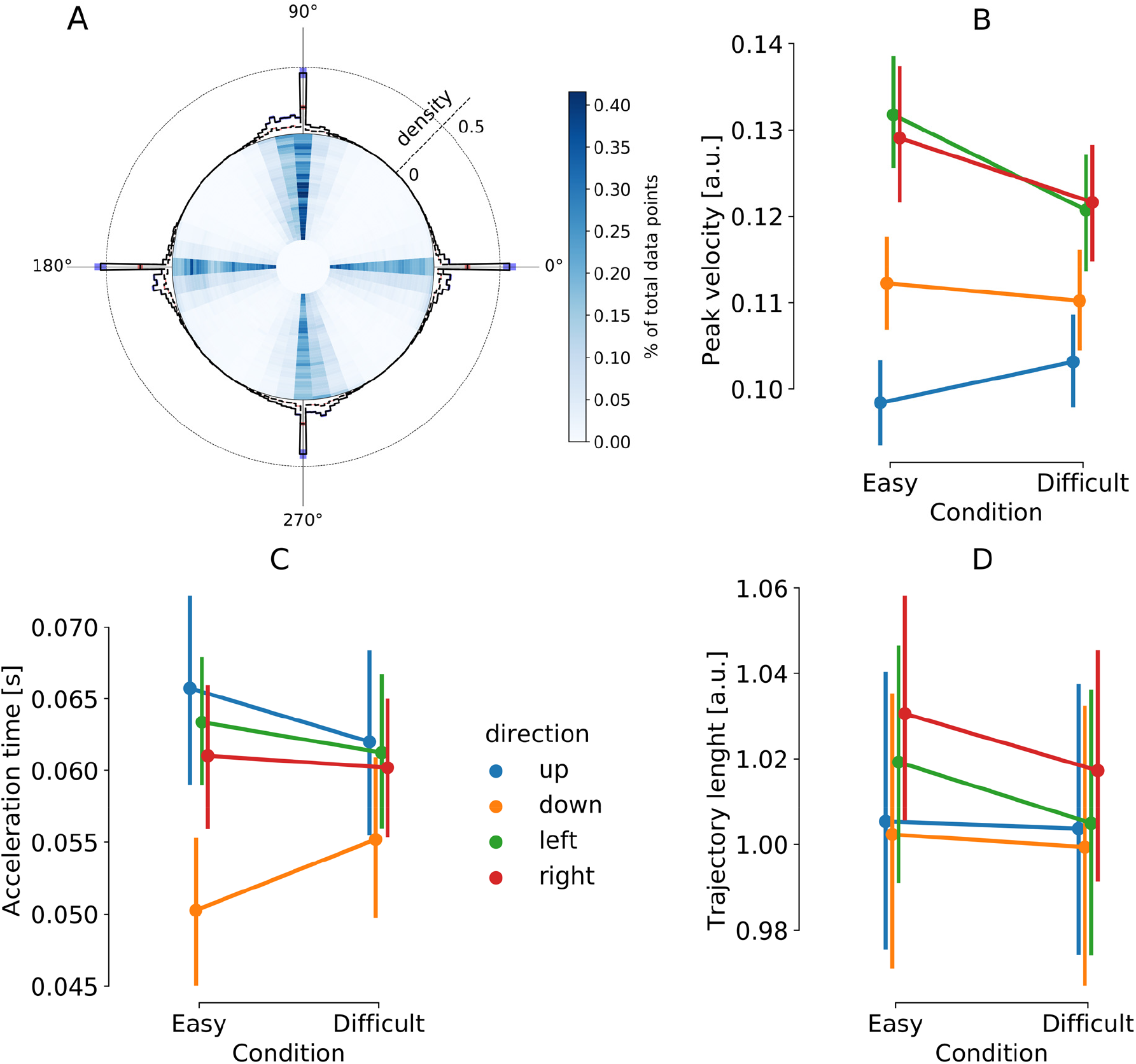
Measures from joystick trajectories. (A) The summary of movement trajectories and final positions. The heat map in the centre represents the proportion of the total joystick position across trials and participants. The histogram on the edge represents the distribution of final positions. (B) The peak velocity of joystick movements averaged across participants. (C) The mean acceleration time of joystick movements averaged across participants (D) The mean trajectory length averaged across participants. The error bars denote the standard errors of the means.

We calculated the action velocity as the rate of changes of joystick position. There was a single peak of action velocity in each trial, consistent with the ballistic nature of the movement. There was strong evidence for the main effect of response direction on the peak velocity (Figure 5B, BF_10_= 3.900 × 10^24^, *F*(2.000, 40.002) = 39.25, *p* < 0.001), moderate evidence for the main effect of difficulty (BF_10_ = 4.612, *F*(1,20) = 22.70, *p* < 0.001) and strong evidence for the interaction between direction and difficulty (BF_10_ = 58.433, *F*(2.841,56.813) = 30.58, *p* < 0.001).

We calculated the acceleration time as the latency between the RT and the time of peak velocity (Figure 5C). There was strong evidence for the main effect of response direction (BF_10_ = 1147.376, *F*(2.253, 45.05) = 4.741, *p* = 0.011). We found moderate evidence against difficulty level (BF_10_ = 0.172, *F*(1,20) =0.178, *p* = 0.677). Frequentist ANOVA showed a significant interaction between the response direction and difficulty levels (*F*(2.853, 57.053) = 4.470, *p* = 0.008), which was not supported by the Bayes factor (BF_10_ = 0.256).

We calculated the trajectory length as the sum of the Euclidean distance between adjacent joystick positions in each trial (Figure 5D). There was no compelling evidence for the main effect of response direction on trajectory length (BF_10_ =1.759; *F*(3, 60) = 1.944, *p* = 0.151), nor the main effect of task difficulty (BF_10_ = 0.450, *F*(1, 20) = 3.171, *p* = 0.09). The evidence against the interaction between direction and difficulty was strong (BF_10_ = 0.090, *F*(3, 60) = 0.978, *p* = 0.409).

In summary, the peak action velocity of joystick movements was affected by both action direction and task difficulty, and acceleration time was affected only by trajectory direction. There was no compelling evidence to support that trajectory length was affected by action direction or task difficulty.

## Discussion

The current study systematically compared the consistency between continuous and discrete responses during rapid decision-making. In a four-alternative motion discrimination task, joystick movements and key presses led to similar accuracy and mean RT. Further modelling analysis with hierarchical DDM showed no evidence in supporting a change of any model parameters between response modalities. Together, our findings provide evidence for the validity of using continuous joystick movement as a reliable response modality in behavioural experiments.

### Behavioural measures

In both joystick and keyboard sessions, participants had lower accuracy and longer mean RT in the more difficult condition (i.e., lower motion coherence), in line with previous findings with similar tasks (Britten et al., 1992; Pilly & Seitz, 2009; Ramachandran & Anstis, 1983; Roitman & Shadlen, 2002). Using Bayesian statistics, we found evidence that response modality (joystick motion or key press) did not affect either accuracy or mean RT, confirming the validity of using joystick as a response device in decision-making tasks. Importantly, across participants, the difference in the RT distributions between response modalities was positively correlated between easy and difficult conditions. Therefore, participants with similar behavioural performance between response modalities maintained their consistency between experimental conditions.

Joystick positions estimated at a high sampling rate enabled additional behavioural measures beyond on/off key presses. In the current study, most of the movement trajectories were along the four cardinal directions (Figure 5A). The averaged trajectory length was close to 1 (Figure 5D), which was the shortest distance from the joystick’s neutral position to the maximum range, suggesting that the participants were able to make accurate and ballistic movements following the task instruction. Nevertheless, it is worth noting that the movement direction affected the peak velocity and acceleration time. This may be due to the difference in upper limb muscle contractions when moving the joystick towards different directions (Oliver, Northey, Murphy, MacLean, & Sexsmith, 2011). Therefore, for future behavioural experiments relying on sensitive trajectories measures, we suggest extra cautious on the effects of ergonomics and human motor physiology, especially for rapid movements as in the current study. One potential solution would be to acquire baseline recordings of the movements to be expected during the experiment, which can then be used to compensate measurement biases.

### Model-based measures

The DDM and other sequential sampling models are commonly used to investigate the cognitive processes underlying rapid decision-making (Bogacz et al., 2006; Smith & Ratcliff, 2004). In the current study, the mean drift rate increased in the easier task condition, consistent with previous modelling results (Ratcliff & McKoon, 2008). The combination of posterior parameter estimation and Bayesian inference allowed us to obtain the probability of the parameter being practically equal, a more informative measure than frequentist *p*-values (Kruschke, 2015). Although our results suggested that most parameter values had high probabilities to remain the same between response modalities (Table 1), we could not accept the null hypothesis for certain (which requires *P*_*P*|*D*_ = 1) and need more data to confirm the inference.

We highlighted two model parameters with low *P*_*P*|*D*_ values, which indicate that, with additional observed data from future experiments, the posterior model parameters might be in favour of the alternative hypothesis (i.e., a difference between response modalities). First, when switching from key presses to joystick movements, there was a small increase in the mean non-decision time (*P*_*P*|*D*_ = 0.658). Second, responding with a joystick resulted in a slightly decreased decision threshold (*P*_*P*|*D*_ = 0.872). Several previous studies showed that instructing to respond faster or more accurately could efficiently modulate participants’ behaviour (Beersma et al., 2003; Schouten & Bekker, 1967; Wickelgren, 1977). The decision threshold plays a substantial role under such speed-accuracy instructions (Mulder et al., 2013; Rae, Heathcote, Donkin, Averell, & Brown, 2014; Ratcliff & McKoon, 2008; Starns & Ratcliff, 2014; Zhang & Rowe, 2014): a decrease of threshold is accompanied with faster reaction speed and lower accuracy. If participants do implicitly trade accuracy for speed when switching from keyboard to joystick movements, this cognitive discrepancy needs to be considered when conducting experiments involving continuous responses. One hypothesis for this potential behavioural change is that continuous joystick movements allow participants to change or correct their responses later in a trial (Albantakis & Deco, 2009; Gallivan & Chapman, 2014; Gallivan, Logan, Wolpert, & Flanagan, 2016; Selen, Shadlen, & Wolpert, 2012), and this response flexibility may lead to reduced deliberation in initial movements.

The trial-by-trial variabilities in drift rate and non-decision time also had *P*_*p*|*D*_ values. Empirically, across-trial variability was introduced in DDM to improve the model fit to RT distributions (Ratcliff & McKoon, 2008), although the functional significance of these parameters to the decision process is still unclear. Across-trial variability in the drift rate produces different RT between correct and error trials (Ratcliff & Rouder, 1998), and across-trial variability in the non-decision time accounts for the large variability in trials with short RT across experimental conditions (Ratcliff & Tuerlinckx, 2002). These model parameters allow the DDM to account for the subtle differences in the shape of RT distributions between response modalities. Future studies could apply formal model comparison to evaluate the need of trial-by-trial variability in modelling joystick responses.

### The use of joystick and its validity

We aimed to establish the validity of joystick response in rapid decision-making tasks. More specifically, we examined whether response modality (joystick movements vs. key presses) alters the raw behavioural measures (RT and accuracy) and underlying cognitive processes. We found that both behavioural measures and model parameters from cognitive modelling did not differ significantly between response modalities. In other words, using joystick movements to indicate choices of perceptual decisions elicit behavioural and cognitive characteristics similar to those from conventional key presses.

Motion discrimination based on random dot kinematogram is a typical paradigm for simple decisions. The same computational mechanism of evidence accumulation has been suggested to account for the cognitive processes underlying a broad range of decision-making tasks, spanning across sensory modalities (O’Connell, Dockree, & Kelly, 2012) and cognitive domains (Gold & Shadlen, 2007). Therefore, we expect that the validity of joystick response established in the current study can be extended to experimental paradigms in which participants make rapid choices with motor actions (Ratcliff & McKoon, 2008).

The joystick as a response modality has been successfully applied in ageing and clinical populations, in which conventional key presses may be error-prone due to impaired dexterity. Both older and young adults can operate joysticks in visuomotor tasks with similar response patterns (Kramer, Larish, Weber, & Bardell, 1999). Previous studies showed that older adults can complete multiple hour-long cognitive training sessions with joystick responses, and the performance benefit persisted for 6 months after training (Anguera et al., 2013). In patients with neurodegenerative diseases, volitional joystick movements have been successfully used to examine the motor deficits and underlying neural abnormalities (Kew et al., 1993). This evidence suggested that the use of joystick can be well tolerated in older adults and patients.

In the current study, the participants did not report fatigue after joystick or keyboard sessions, which lasted approximately 45 minutes each. Other paradigms with longer experimental sessions and more intense joystick movements may impose a challenge to participants’ stamina. Nevertheless, it is possible to use measures from the continuous joystick recording (Kahol, Smith, Brandenberger, Ashby, & Ferrara, 2011) or concurrent physiological recording (Mascord & Heath, 1992) to identify the onset of fatigue prior to performance deterioration.

One may ask if joystick responses provide any additional value over conventional key presses. Here, we showed that, even in simple ballistic movements, joystick-specific measures (e.g. action velocity) can be affected by the task difficulty, providing additional information on behavioural performance in addition to RT and accuracy. It is yet to be determined whether continuous responses provide independent information from discrete responses (Freeman, 2018; Freeman & Ambady, 2010; Stillman, Medvedev, & Ferguson, 2017). However, the capacity of recoding continuous responses via joysticks enables new experimental designs to probe the continuous interplay between action, perception and cognition. For example, the ongoing locomotion can modify the sensory information flow (Ayaz, Saleem, Schölvinck, & Carandini, 2013; Souman, Freeman, Eikmeier, & Ernst, 2010).

### Future directions

Three issues require further consideration. First, we only used a joystick to record movement trajectories, which is commonly used and widely available in behavioural testing labs. There are many other devices capable for recording continuous responses, such as computer mouse (e.g. Koop & Johnson, 2011), optic motion sensor (e.g. Chapman et al., 2010) and robotic arms (Abrams, Meyer, & Kornblum, 1990; Archambault, Caminiti, & Battaglia-Mayer, 2009; Berg et al., 2016; Burk, Ingram, Franklin, Shadlen, & Wolpert, 2014; Resulaj et al., 2009). The current study offered a comprehensive comparison between key presses and joystick movements, but the measures from other devices are yet to be validated. We also offered a practical solution to measure RT from joystick movement comparable to that from key presses, taking in to account the small resistive forces near the joystick’s neural position. To facilitate future research, we have made our data and analysis scripts openly available (https://osf.io/6fpq4).

Second, we instructed participants to make directional movements in the joystick session, which allows for intra-individual comparisons between response modalities. Motion trajectories suggested that the participants mainly made ballistic actions towards one of the four cardinal directions (Figure 5A). One could explore the further potential of continuous responses in behavioural tasks, such as in response to the change of mind (Berg et al., 2016; Burk et al., 2014; Resulaj et al., 2009) or external distractions (Gallivan & Chapman, 2014).

Third, the DDM required the behavioural data to be presented as binary choices (Ratcliff & McKoon, 2008). To meet this constraint, we simplified our four-choice task data into correct and incorrect decisions, and incorrect responses contained errors towards three different directions from the correct motion direction. Our modelling results provided a good fit to the observed data. It would be useful to extend the analysis using other models that are designed for decision problems with multiple alternatives (Bogacz, Usher, Zhang, & McClelland, 2007; Brown & Heathcote, 2008; Usher & McClelland, 2001; Wong & Wang, 2006; Zhang & Bogacz, 2009), although a hierarchical Bayesian implementation of those more complex models is beyond the scope of the current study.

In conclusion, our results validated the joystick as a reliable device for continuous responses during rapid decision-making. Compared with key presses, the additional complexity and continuity associated with joystick movements did not affect raw behavioural measures such as accuracy and mean RT, as well as underlying decision-making processes. However, we highlighted the effects of movement direction on continuous trajectory measures. Researchers should be cautious when adopting experimental designs that require complex movement trajectories.

## Acknowledgements

MJS was supported by a PhD studentship from Cardiff University School of Psychology. JZ was supported by a European Research Council Starting grant (716321). The authors declare no competing financial interests. We want to thank Simon Rushton for the helpful comments.

## Open practices statement

All the data and the materials for the experiment and analysis are available at https://osf.io/6fpq4

---

**Supplementary Figure 1.**
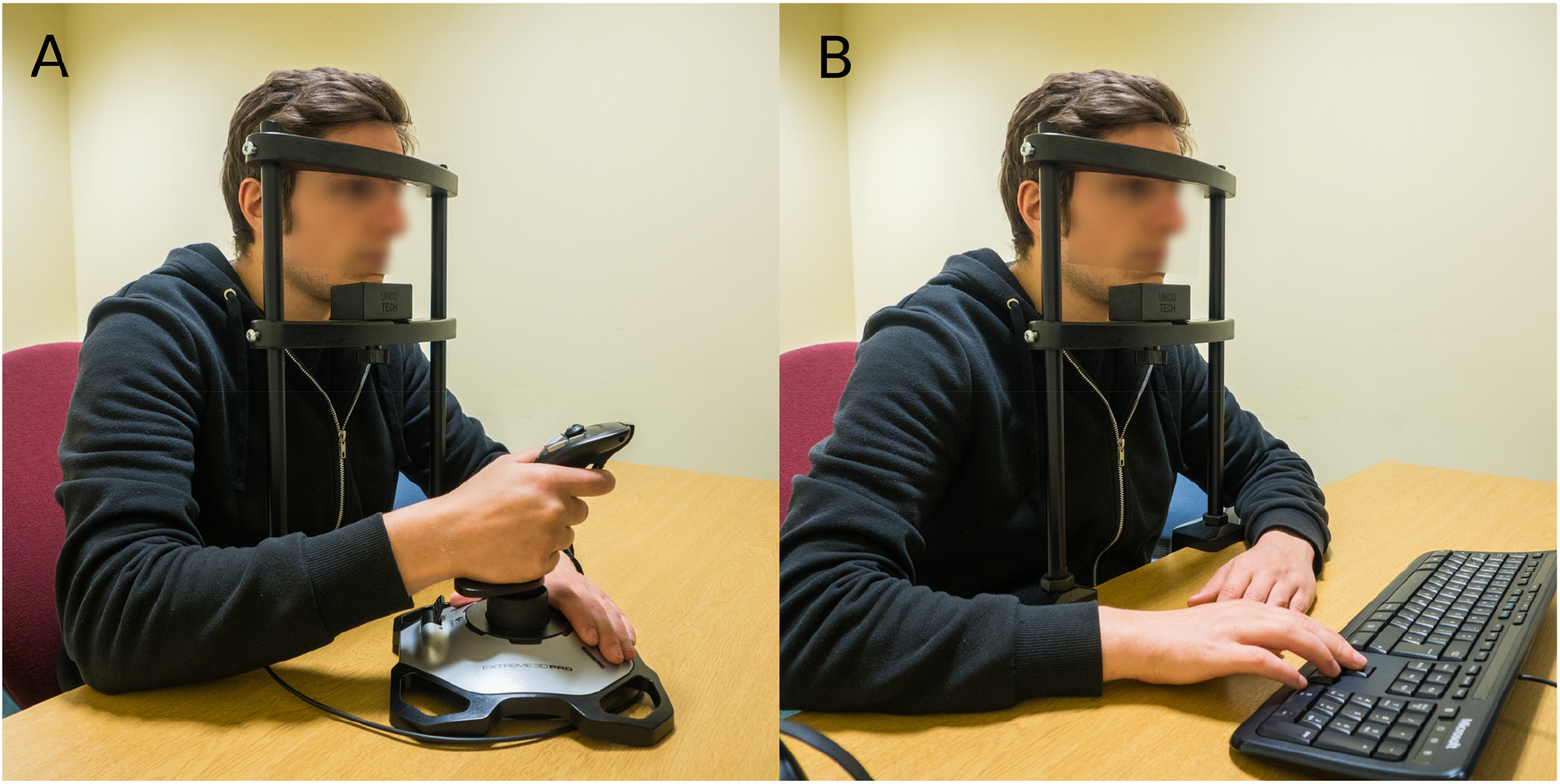
The experimental setup and joystick positioning. Participant was seated in front of the screen. The distance from the screen and the head position was maintained using a chinrest. Seating height was adjusted to the most comfortable position. Joystick was positioned to the right of the participant (A). Exact position of the device was adjusted to the most comfortable position. Participants were asked to hold the base of the joystick while responding. Keyboard was placed parallel to the screen to ensure the arrow directions correspond to the direction of the motion of the visual stimuli (B).

